# MSA: Reproducible mutational signature attribution with confidence based on simulations

**DOI:** 10.1101/2020.12.14.422764

**Authors:** Sergey Senkin

## Abstract

**Background:** Mutational signatures proved to be a useful tool for identifying patterns of mutations in genomes, often providing valuable insights about mutagenic processes or normal DNA damage. *De novo* extraction of signatures is commonly performed using Non-Negative Matrix Factorisation (NMF) methods, however, accurate attribution of these signatures to individual samples is a distinct problem requiring uncertainty estimation, particularly in noisy scenarios or when the acting signatures have similar shapes. Whilst many packages for signature attribution exist, a few provide accuracy measures, and most are not easily reproducible nor scalable in high-performance computing environments.

**Results:** We present MSA (Mutational Signature Attribution), a reproducible pipeline designed to assign signatures of different mutation types on a single-sample basis, based on Non-Negative Least Squares (NNLS) method with optimisation. Parametric bootstrap is proposed as a way to measure statistical uncertainties of signature attribution. Supported mutation types include single and doublet base substitutions, indels and structural variants. Results are validated using simulations with reference COSMIC signatures, as well as randomly generated signatures.

**Availability and implementation:** MSA comprises a set of Python scripts unified in a single Nextflow pipeline with containerisation for cross-platform reproducibility and scalability in high-performance computing environments. The tool is publicly available from https://gitlab.com/s.senkin/MSA

## Background

Mutational signatures are distinctive combinations of somatic mutations which can be of various origin, such as exogenous or endogenous exposures, defective DNA repair pathways, DNA replication infidelity or DNA enzymatic editing [1, 2]. Currently on the order of 100 signatures have been discovered in human cancer, but for the majority of them the aetiology remains unknown. This has given rise to a rich new field of mutational signature discovery, as well as linking signatures to various risk factors, in projects like Mutographs [REF].

*De novo* extraction of signatures is commonly performed using Non-Negative Matrix Factorisation (NMF)[3] for somatic mutations under various mutational classifications[4], with tools such as SigProfilerExtractor [REF]. Such signature extraction has been extremely informative in the analysis of many cancer types and shed light into mutagenesis of endogenous and exogenous risk factors[2].

It has also become apparent that certain signatures show a dose-response relationship with risk factors, for example COSMIC signature SBS4 with tobacco smoking[5]. In order to characterise such relationships, it is increasingly important to attribute mutational signatures, i.e. estimate their activities in any given sample, with confidence intervals. Some signatures with similar shapes are difficult to differentiate between each other, for instance COSMIC signatures SBS5 and SBS40 that both have a relatively flat profile. For such signatures, point estimates of attribution can often be inaccurate, leading to false positive or false negative findings.

Ideally, statistical uncertainty of signature attribution would be best estimated by performing repeated measurements. Given the high cost and complexity of such measurements (especially sample preparation, DNA extraction and sequencing), one needs to look for other alternatives. Bootstrapping has been proposed as a practical method to estimate uncertainty of signature attribution[6, 7], however, not all investigators are explicit in the precise version of bootstrapping used. Whereas some suggest that simple resampling with replacement of a mutationol catalogue can give a meaningful result, we argue that classic bootstrap is not applicable in mutational signature attribution, and propose the parametric bootstrap approach under assumption that mutations accumulate according to Poisson processes.

Another common difficulty of signature attribution packages is the low reproducibility as well as scalability of computations in high-performance computing environments. This is a particular concern when analysis is performed on large datasets, and when the number of bootstrap variations is considerable. We resolve this issue by utilising Nextflow[8], a domain-specific language (DSL) designed to primarily address computational irreproducibility and efficient parallel execution in a large number of computing environments, from individual workstations to server clusters and cloud computing services.

## Approach

Consider a mutational catalogue (such as a set of somatic mutations in cancer genomes) as a matrix **M**(*m× n*). Here, *m* is the number of mutation types, such as 96 single base substitution types in the trinucleotide context, and *n* is the number of samples. Let **S**(*m × k*) be the non-negative matrix of *k* mutational signatures (e.g. COSMIC catalogue[2]). The goal is to find the non-negative matrix of activities (or exposures) **A**(*k × n*):

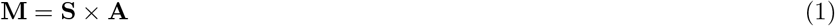

In a more general version of the problem where the matrix **S** is unknown, *de novo* extraction of signatures can be performed using algorithms such as Non-negative Matrix Factorisation (NMF). Here, we assume that the signature matrix **S** is known, thereby scrutinising a more specific problem of finding activities of predefined signatures. This is particularly relevant when estimating the uncertainty of each signature activity is important, or simply when the number of samples *n* is low and one needs to quickly examine how well these samples can be explained by known mutational signatures.

The signature attribution problem can therefore be studied independently for each given sample **y** from the mutational catalogue **M**, as it can be seen as a linear combination of signatures **S** and their per-sample activities **x**:

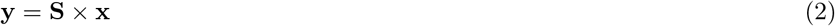

Several algorithms can be used to solve this kind of problem, such as quadratic programming (QP) or simulated annealing (SA). Here, we utilise the Lawson-Hanson algorithm for non-negative least squares (NNLS) [9], which is essentially a constrained version of the ordinary least squares problem:

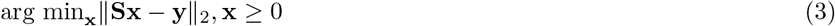

Application of NNLS out of the box can be performed using one of the popular packages (*nnls* in R or *optimize.nnls* function in *scipy* library in Python), however, such approach is known to over-fit the data and lead to a large number of false-positive findings, as shown using simulations below. To mitigate this, ad-hoc approaches can be utilised, such as the optimisation of signature attribution using additional penalties on each signature’s contribution, or by using pre-existing biological knowledge such as strand bias rules [2, 10]. Although such optimisation is implemented in MSA using the penalty loops described below, the main advantage of the tool is the ability to quantify the confidence of attribution for each signature.

### Classic bootstrap

To estimate the confidence of signature attribution, several bootstrap approaches have been explored when developing this tool. In principle, bootstrapping can be applied to the full genome sequence data, but in the context of mutational signatures we apply bootstrap to collapsed mutational catalogues (matrix **M** in Equation (1)) for the sake of computational facility. Furthermore, it appears problematic to validate bootstrapped full sequences with simulations of predefined signatures.

Initially, a simple bootstrap, meaning resampling with replacement, was attempted on both simulated and real data. In this approach, *N* observations are randomly drawn from an initial sample with *N* mutation counts with replacement. In any given resample, each observation occurs 0, 1 or more times according to the binomial distribution *Binomial*(*N,* 1*/N*). Since the total number of observations is *N*, all counts jointly form a multinomial distribution *Multinomial*(*N,* 1*/N, …,* 1*/N*). Importantly, this approach gives meaningful results only under the i.i.d. assumption, i.e. when underlying random variables are independent and identically distributed[11]. This is not the case in mutation profiles: existing mutational signatures suggest that accumulation of mutations in particular trinucleotide contexts are inter-dependent and not identically distributed. For example, samples with high levels of APOBEC activity are characterised by mutations enriched within TCA and TCT contexts, clearly not identically distributed to all other trinucleotide contexts. Not surprisingly, using simple bootstrap to derive confidence intervals of signature attribution leads to rather poor results (Figure 1 (a)). In a simulated example, four COSMIC signatures (SBS 1, 5, 22 and 40) were used to mimic real kidney RCC data without noise. Whilst running NNLS on the input sample yielded attributions nearly identical to the simulated true values, results on bootstrapped samples were vastly different, producing grossly inaccurate confidence intervals.

**Figure 1.**
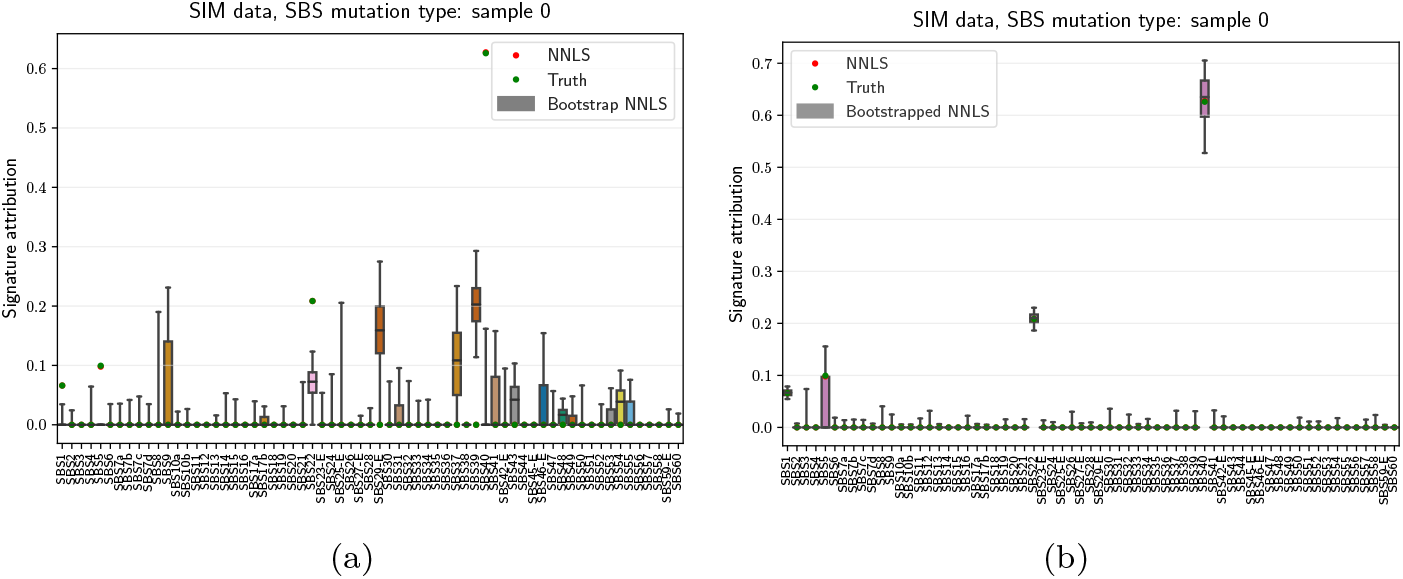
Attribution with (a) classic and (b) parametric bootstrap. Confidence intervals are derived as [2.5%, 97.5%] percentiles of the resulting bootstrap activities, applied to simulated data with COSMIC signatures SBS 1, 5, 22 and 40, without noise. Point estimates of NNLS attribution (red dots) almost perfectly coincide with the generated true values (green dots) in such noiseless scenario.

Some investigators argue that classic bootstrap approach can be regarded as conservative for non-i.i.d. data[12]. Other investigators argue that classic bootstrap is inadequate for high-dimensional non-i.i.d. data[13]. In the context of highdimensional mutational profiles, we choose to derive confidence intervals on signature attributions using parametric bootstrap.

### Parametric bootstrap using Multinomial distribution

Parametric bootstrap assumes that the data follows a known underlying distribution. This implies making certain assumptions on the original dataset, so that the bootstrap samples can be drawn from the estimated parametric model.

Here, we make an assumption that mutations are accumulated following Poisson distributions for each mutation type, i.e. that mutations accumulate randomly, independently and at a constant rate.

The idea of parametric bootstrap applied to mutational processes was inspired by *mutSignatures* package[7] (which is itself based on the original MATLAB framework for deciphering signatures[14]), where the input mutation matrix is bootstrapped according to the multinomial distribution *Multinomial*(*M, p*_1_*, …, p_m_*), where *M* is the total mutational burden in a given sample, and probabilities *p_i_* are normalised mutation counts for each mutation type. This distribution is chosen since the conditional distribution of a vector of independent Poisson variables is equivalent to multinomial distribution[15].

Since the mutational burden of each bootstrap samples is fixed and equal to *M*, we slightly modify the method by drawing counts from independent binomial distributions, so that the total mutational burden is no longer fixed. Nevertheless, for any given mutation category (e.g. *C*[*T* > *A*]*G* trinucleotide context on Figure 2) the distribution of bootstrapped mutation counts follows a Poisson distribution.

**Figure 2.**
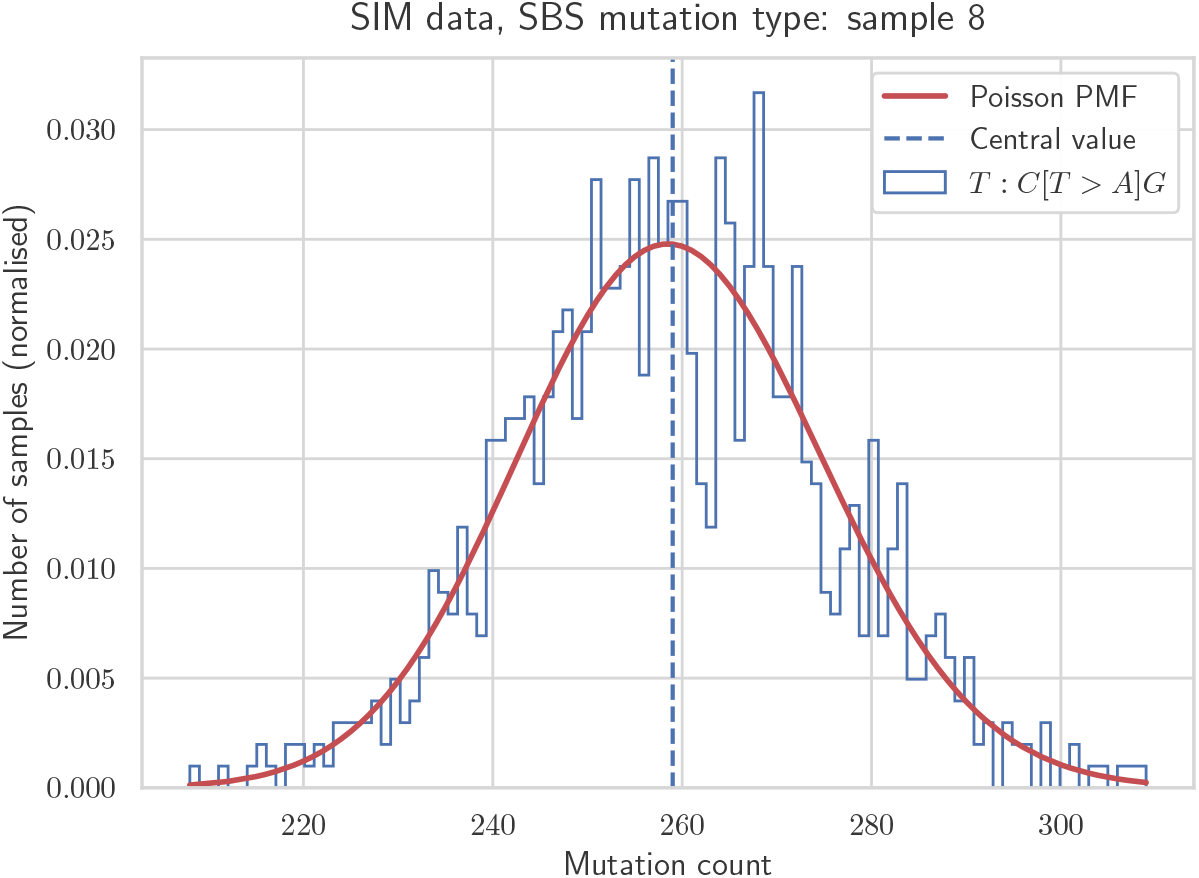
Example of parametric bootstrap distribution for a *C*[*T* > *A*]*G* trinucleotide context. Simulated sample with 1000 bootstrap variations, the histogram is fitted by Poisson probability mass function.

For each bootstrap sample, NNLS attribution is applied to derive the vector of signature activities. 95% confidence intervals are then derived for each signature attribution by taking [2.5%, 97.5%] percentiles of the resulting bootstrap activities. Direct comparison with classic bootstrap on a simple simulated case without noise shows a clear advantage of the parametric method (Figure 1 (b)).

### Optimisation of signature attribution

Since pure NNLS is based on a simple fitting approach, it is generally prone to over-fitting, particularly in noisy environments[16]. To mitigate this, a form of regularisation can be applied using the penalisation loops to add or remove signatures based on their contribution to the fit. This approach is based on the penalised attribution used in PCAWG consortium signature assignment[2].

The default optimisation strategy, called removal strategy, starts with a full set of available signatures. Base *L*2 similarity of the reconstructed profile (linear combination of input signatures) to the input mutational profile is calculated, normalised by its *L*2 norm. Afterwards, a removal loop is executed, where all least contributing signatures increasing the *L*2 similarity by less than a given penalty (called ‘weak” threshold) are sequentially removed. The resulting set of remaining signatures is used to describe the input sample by applying the final NNLS fit.

On the other hand, high penalties used in optimisation can lead to under-fitting of data, meaning that optimal penalties need to be derived. We perform this by running simulations and measuring sensitivities, specificities as well as other metrics for all acting signatures.

### Validation with simulations

When simulating a dataset that is supposed to resemble real data, one generally has to make certain assumptions about underlying distributions. Firstly, we explored scenarios mimicking real cohorts such as RCC (shown on Figure 1), with a defined set of acting reference COSMIC signatures, where generated signature attributions follow non-negative zero-inflated Gaussian distributions. Generally, such distributions do not describe real attributions well, especially for rare signatures. Therefore, as a second approach, we explored simulations based on a simple bootstrap of signature activities derived with very low penalties, where such activities are used to generate mutational profiles which are then resampled with replacement. These activities are bound to be over-fitted, yet they provide a way to sample signature attributions from distributions of observed real data. The over-fitted generated signatures generally have low attribution levels, and can be regarded as noise in addition to the default noise model. We use the Poisson noise model by default, where the variance is equal to the mean generated mutation burden for any given mutation type.

Simulations allow to estimate the performance of signature attribution overall, as well as for each signature in different optimisation scenarios. Since the simulated truth is always known, one is able to calculate various metrics in order to estimate the accuracy of signature attribution, such as sensitivity and specificity for each signature, or for all signatures on average. Figure 3 shows such metrics for all COSMIC SBS signatures combined, based on the simulated model of ESCC signature activities derived with the *L*2 penalty of 0.001. Figure 4 shows these metrics for COSMIC signatures SBS5 and SBS16, for a range of “weak” penalties discussed previously. Signatures SBS5 and SBS16 were picked as typical examles of signatures with flat and non-flat profiles, respectively. The L2 penalty of 0 corresponds to simple NNLS fit without any optimisation. Metrics are estimated with and without utilising confidence intervals, with the latter approach using the lower limit of CIs when calculating metrics.

**Figure 3.**
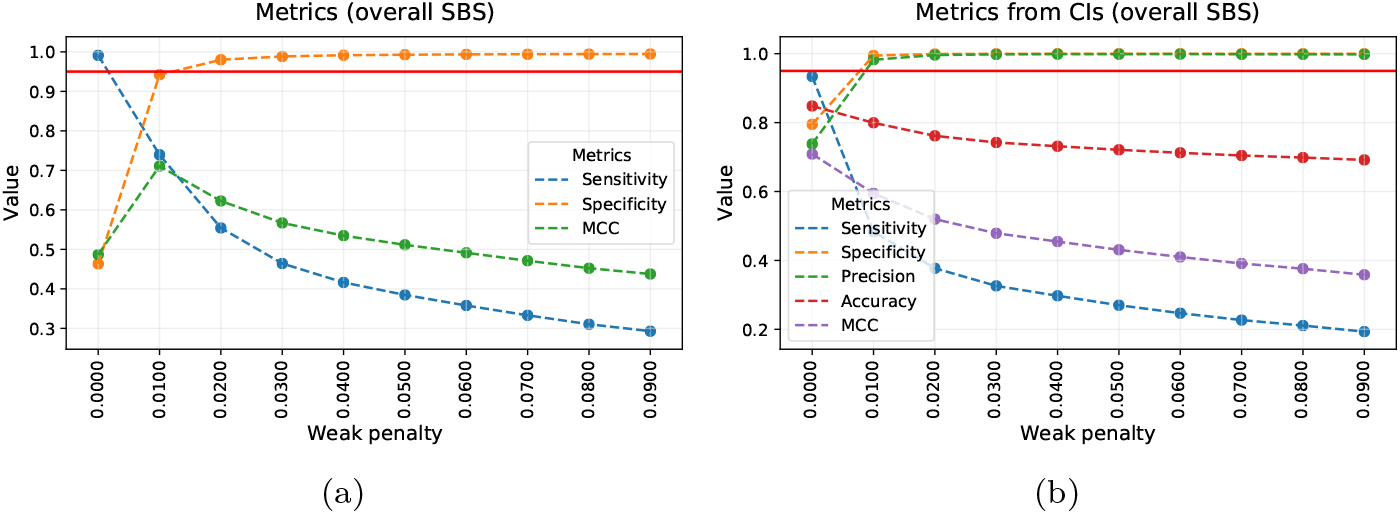
Overall performance metrics of signature attribution for simulated ESCC cohort with respect to optimisation penalty. (a) Senstivity, specificity and Matthews correlation coefficient are calculated using point estimates without confidence intervals. (b) Similar metrics are calculated using bootstrap variations yielding confidence intervals. Red horizontal line corresponds to the 95% value for each metric.

**Figure 4.**
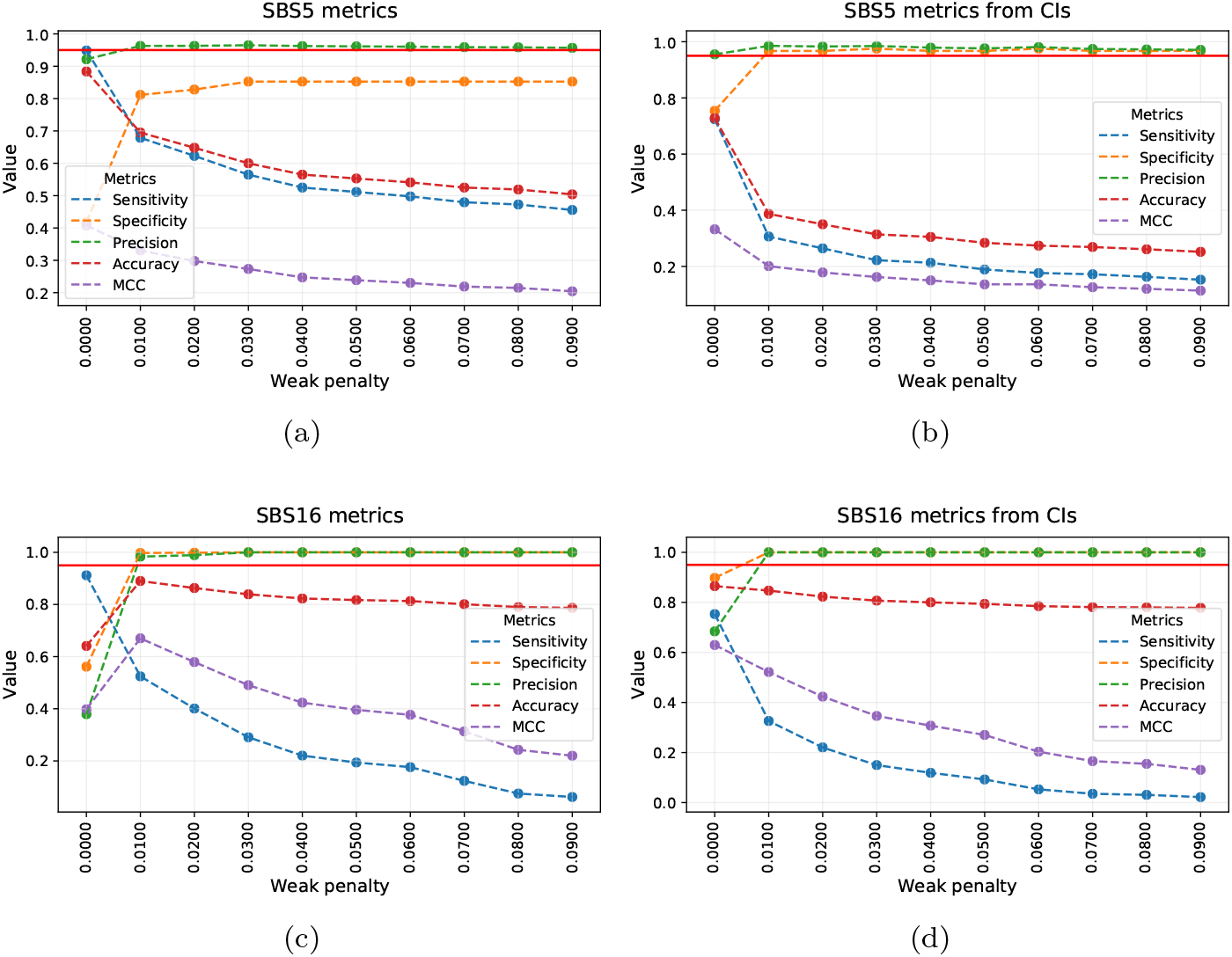
Performance metrics for COSMIC signatures SBS5 and SBS16 with respect to optimisation penalty. SBS5 metrics are derived using point estimates (a) and confidence intervals (b). Similarly, SBS16 metrics are derived using point estimates (c) and confidence intervals (d). Red horizontal line corresponds to the 95% value for each metric.

First of all, it is evident that higher penalties lead to lower sensitivity yet higher specificity of signature attribution. Secondly, confidence intervals allow to reach higher levels of specificity for signature SBS5 that is particularly difficult to attribute due to its relatively flat profile, and similarity to another acting signature, SBS40. Without confidence intervals, i.e. only using point estimates of attributions derived for simulated samples, specificity of SBS5 never reaches 90%. On the other hand, confidence intervals do drive weakly-acting signatures with low attributions to zero, hence lowering the overall sensitivity to such signatures. Generally, finding an optimal penalty is always a trade-off, and investigators are free to choose a more or less conservative path by tweaking such parameters.

## Implementation

The MSA tool is implemented as a set of scripts in Python language, fully automised using Nextflow workflow management system[8]. All dependencies, including packages such as *pandas, numpy, scipy, matplotlib* and *seaborn*, are automatically handled by Nextflow via containerisation using Docker technology, as well as Singularity where Docker is not available. Users not willing to use Docker or Singularity may opt to use a Conda environment that is also automatically handled by Nextflow, yet this approach has inferior reproducibility compared to container technology.

Native support of SigProfilerMatrixGenerator[4] and SigProfilerExtractor [REF] tools is implemented in MSA for convenience. Where SigProfiler outputs are available, running MSA is as simple as executing a single line containing the output paths:

~~~
nextflow run https://gitlab.com/s.senkin/MSA -profile docker
--dataset test --SP_matrix_generator_output_path path/to/SP_ME/
--SP_extractor_output_path path/to/SP_extractor/
~~~

Mutational classifications currently supported are SBS-96 and SBS-288 for single base substitutions signatures, DBS-78 for doublet base substitutions signautres, ID-83 for small insertion and deletion signatures, SV-32 for structural variants signatures.

## Discussion

We explored parametric bootstrap based on multinomial distribution as an effective method to derive confidence intervals of mutational signature attribution for any given mutational profile and available set of *de novo* extracted, or reference signatures. The multinomial distribution is chosen as the one corresponding to an assumption that mutations are accumulated according to Poisson processes for each mutation type, but other assumptions can be potentially investigated – such as Monte Carlo simulations of mutations following distributions of increasing complexity.

The main limitation of the parametric bootstrap we use is its bias towards observed data, since empirical probabilities *p_i_* in the multinomial distribution *Multinomial*(*N, p_1_, …, p_m_*), are taken from real data which can be inaccurate. However, this method aims to estimate the statistical uncertainty of signature attribution method rather than total uncertainty of attribution, and for such purposes remains adequate.

Parametric bootstrap can be applied using both simple NNLS attribution and the one based on penalised optimisation designed to maximise sensitivity and specificity of signature attribution. We developed a set of tools assisting in the validation of optimisation parameters for any given scenario using simulations of different cancer types and population cohorts. In general, it appears that such optimisation is unique for any given cohort due to the uniqueness of acting signatures. It also depends on noise, therefore investigators exploring cohorts of interest ideally need to test different noise scenarios. We opted to pick the Poisson noise model, yet other models such as negative binomial noise are worth consideration.

Estimating the systematic uncertainties of signature attribution, such as the ones due to sequencing artefacts, is a more challenging task that requires comparison of multiple sequencing technologies, not feasible in most settings. However, an uncertainty due to the variant calling can potentially be estimated for any given variant, and propagated to the signature attribution uncertainty. Evaluating these and other uncertainties remains a matter for further studies.

In summary, considering different sources of errors is an important exercise for any measurement, and as we show here, particularly for the attribution of mutational signatures. We hope that investigators will continue to ask such questions and thrive to advance the methods shedding light on them. As a step in this direction, we present MSA – a computational tool to attribute mutational signatures with confidence intervals in an easily reproducible and scalable manner.

## Availability and requirements

- **Project name**: MSA
- **Project home page**: https://gitlab.com/s.senkin/MSA
- **Operating system(s)**: Platform independent
- **Programming language**: Python, Nextflow DSL
- **Other requirements**: Java 8 or later, docker, conda or singularity
- **License**: GNU GPL
- **Any restrictions to use by non-academics**: None

## Acknowledgements

This work was supported by a Cancer Grand Challenges Mutographs team award funded by Cancer Research UK [C98/A24032]. The funder had no role in study design, data collection and analysis, decision to publish, or preparation of the manuscript. We would like to thank Drs Ludmil Alexandrov, Graham Byrnes, Liacine Bouaoun, Vivian Viallon and Karl Smith-Byrne for immensely useful discussions and feedback.

## Disclaimer

Where authors are identified as personnel of the International Agency for Research on Cancer / World Health Organization, the authors alone are responsible for the views expressed in this article and they do not necessarily represent the decisions, policy or views of the International Agency for Research on Cancer / World Health Organization.

